# Challenges in identifying mRNA transcript starts and ends from long-read sequencing data

**DOI:** 10.1101/2023.07.26.550536

**Authors:** Ezequiel Calvo-Roitberg, Rachel F. Daniels, Athma A. Pai

## Abstract

Long-read sequencing (LRS) technologies have the potential to revolutionize scientific discoveries in RNA biology, especially by enabling the comprehensive identification and quantification of full length mRNA isoforms. However, inherently high error rates make the analysis of long-read sequencing data challenging. While these error rates have been characterized for sequence and splice site identification, it is still unclear how accurately LRS reads represent transcript start and end sites. Here, we systematically assess the variability and accuracy of mRNA terminal ends identified by LRS reads across multiple sequencing platforms. We find substantial inconsistencies in both the start and end coordinates of LRS reads spanning a gene, such that LRS reads often fail to accurately recapitulate annotated or empirically derived terminal ends of mRNA molecules. To address this challenge, we introduce an approach to condition reads based on empirically derived terminal ends and identified a subset of reads that are more likely to represent full-length transcripts. Our approach can improve transcriptome analyses by enhancing the fidelity of transcript terminal end identification, but may result in lower power to quantify genes or discover novel isoforms. Thus, it is necessary to be cautious when selecting sequencing approaches and/or interpreting data from long-read RNA sequencing.

## INTRODUCTION

The development of long-read sequencing (LRS) technologies has heralded a new generation of mRNA characterization, including the ability to interrogate RNA at single-molecule resolution. These advances move beyond single (local) exon analysis and allow the annotation of mRNA isoforms to inform the full composition of protein coding exons on individual mRNAs (1–3). LRS can also be used to determine the connections between protein coding exons and untranslated regions (UTRs) that regulate mRNA export, localization, translation, and decay (4). Finally, the quantification of mRNA isoforms using LRS can provide data on their relative usage across cell types (5–8), cellular contexts (9), and disease conditions (10), which is important for research related to both disease mechanisms and public health (11, 12). Together, LRS has the potential to enable a functional understanding of how and why specific full length mRNA isoforms are expressed.

Despite great promise, there remain challenges in the widespread implementation of long-read technologies for RNA-based applications. These challenges include relatively low genomic coverage (compared to short-read technologies), which limits the annotation or quantification of lowly-expressed genes or isoforms (13); and persistent and high error rates due to insertion, deletion, or misalignment errors, leading to problems with the accurate identification of single-nucleotide variations (SNVs) and internal splice sites (14–17).

Moreover, although much focus has been placed on the advantages and challenges of LRS in defining exon composition and structure in mRNA isoforms, less careful characterization has been done on the ability to identify and quantify the terminal ends of isoforms - specifically, the transcription start and end sites. Such characterization is crucial for completely delineating the full mRNA molecule and regulatory functions encoded in UTRs. The evaluation of the terminal ends of RNA molecules is impacted by the same challenges described above. In particular, the low accuracy of LRS reads may affect the ability to identify precise sites of transcription initiation or termination (which are not as delineated by sequence elements as splice sites). Although analytical tools have been proposed to correct for these challenges in other parts of RNA molecules (16, 18, 19), it remains unclear how these and/or other technical challenges may uniquely impact terminal end characterization. While previous studies have suggested the possibility of truncation at terminal ends of RNA reads (especially the 5’ end) (20–22), to our knowledge, none have systematically characterized these sources of error, including (a) to what extent does truncation occur at the 5’ and 3’ ends of LRS RNA reads?, (b) how does such truncation affect the accurate determination of RNA terminal ends?, and (c) do reads with incorrect terminal ends impact expression quantification?

To address this gap, we analyzed previously generated LRS data readily available in the public domain to characterize the terminal end identification and quantification, using data produced by two LRS technological platforms; Oxford Nanopore Technologies (ONT), and Pacific Biosciences (PacBio). We find widespread truncation of LRS reads at the 5’ end of mRNA molecules and our analyses reveal critical factors that must be considered when applying LRS technologies to specific biological questions about mRNA isoform usage.

## METHODS

### Downloading sequencing data

Processed TIF-seq2 data containing the coordinates and usage of transcription start-end pairs for two replicates from K562 cells were downloaded from the Gene Expression Omnibus (GSE140912). Fastq files from Oxford Nanopore directcDNA and directRNA reads were downloaded from the Singapore Nanopore-Expression Project (SG-NEx) website (https://github.com/GoekeLab/sg-nex-data) (23). Fastq files from Pacific Biosciences Iso-Seq reads were downloaded from the ENCODE Project data portal (https://www.encodeproject.org/) (24). The specific samples and accession numbers can be found in Table S1.

### Long-read data mapping and processing

All long-read sequencing data were mapped to the hg38 human reference genome (25) using minimap2 (26) following the developers’ recommendations. Specifically, for each sequencing type, the -ax option was modified as follows: *-ax splice* for directcDNA, *-ax splice -uf -k14* for directRNA, *-ax splice:hq -uf* for Iso-seq. To assign reads to genes, primary alignments were split into individual features using *bedtools bamtobed* and then *bedtools intersect* was used to intersect the features with annotated genes. Only the reads where the start and end features were assigned to the FE and polyA peaks for the same gene, respectively, were considered and used in the analysis of the terminal site. The length of the primary aligned reads was calculated from the bam files. Counts per million were calculated by dividing the count of each gene by the total counts in the sample and multiplying by one million.

### Terminal end classification and filtering

We used databases of empirically derived 5’ end start sites or 3’ end polyA sites to assess the accuracy of terminal ends in LRS reads. Specifically, we used two 5’ start site annotations: (a) human CAGE peaks, downloaded from the FANTOM project website (27) and (b) human HITindex first exons (28). HITindex first exons were obtained by running the HITindex pipeline on short-read RNA-seq data from the entire GTeX V8 database (29) and retrieving exons identified as first exons or hybrid first-internal exons in at least one tissue. For 3’ end annotations, we used experimentally verified human polyadenylation sites from the PolyASite 2.0 database (30), which is a consolidated atlas of polyadenylation sites from publicly available 3’ end sequencing datasets.

LRS reads were classified by their overlap with 5’ end start sites and/or 3’ end polyA sites from the sources above. For start sites, we intersected the upstream most feature of each read with either CAGE peaks or HITindex first exons using bedtools intersect and considered overlap as read features that overlapped within +/- 75nt from the CAGE peak or exon 5’ and 3’ ends. Similarly, for end sites, we intersected the downstream most feature of read with polyA peaks from the PolyASite database using bedtools intersect and considered overlap as read features that overlapped within +/- 50nt of the peak 5’ and 3’ ends.

To identify reads most likely to have accurate terminal ends, we retained reads whose terminal features overlapped both a HITindex first exon and a polyA peak from the same gene. The high number of single feature (exon) reads that matched both a CAGE peak and polyA peak within the same exon led us to perform filtering only with HITindex first exons. For all these analyses we also discarded reads that started and ended in the same exon.

## RESULTS

We set out to systematically assess the relative accuracy of transcript start and end sites from LRS RNA approaches. To do so, we downloaded LRS data for direct RNA (ONT), direct cDNA (ONT), and Iso-Seq (PacBio) approaches, each of which have different RNA and library preparation steps prior to sequencing (**Table S1**). Specifically, direct RNA from ONT sequences native RNA from total or enriched samples following the optional synthesis of the complementary strand. The direct cDNA method from ONT uses a strand-switching protocol to prepare full-length cDNAs from polyA+ RNA. Finally, Iso-Seq from PacBio also performs strand-switching to produce a SMRTbellTM template, which are double-stranded DNA templates capped with a hairpin loop (blunt adapters) on each end to generate sense and antisense sequences from a single molecule of cDNA. This structure facilitates error correction using the company’s circular consensus sequencing approach. We also used data from TIF-Seq2 - a protocol that identifies connections between transcript start and end sites on the same short sequencing read - as an orthogonal, non-LRS control (31). All LRS data was mapped to the human genome and processed in a similar manner to ensure comparability (Methods). Using these data, we identified and compared transcript start and end sites across sequencing technologies.

We find that there is a substantial heterogeneity in isoforms identified for a same gene using LRS reads from different approaches. Two primary patterns stood out and are exemplified in the representative visualization of reads from the CDC42 gene using LRS and TIF-Seq2 data from K562 cells (**Figure 1**).

**FIGURE 1:**
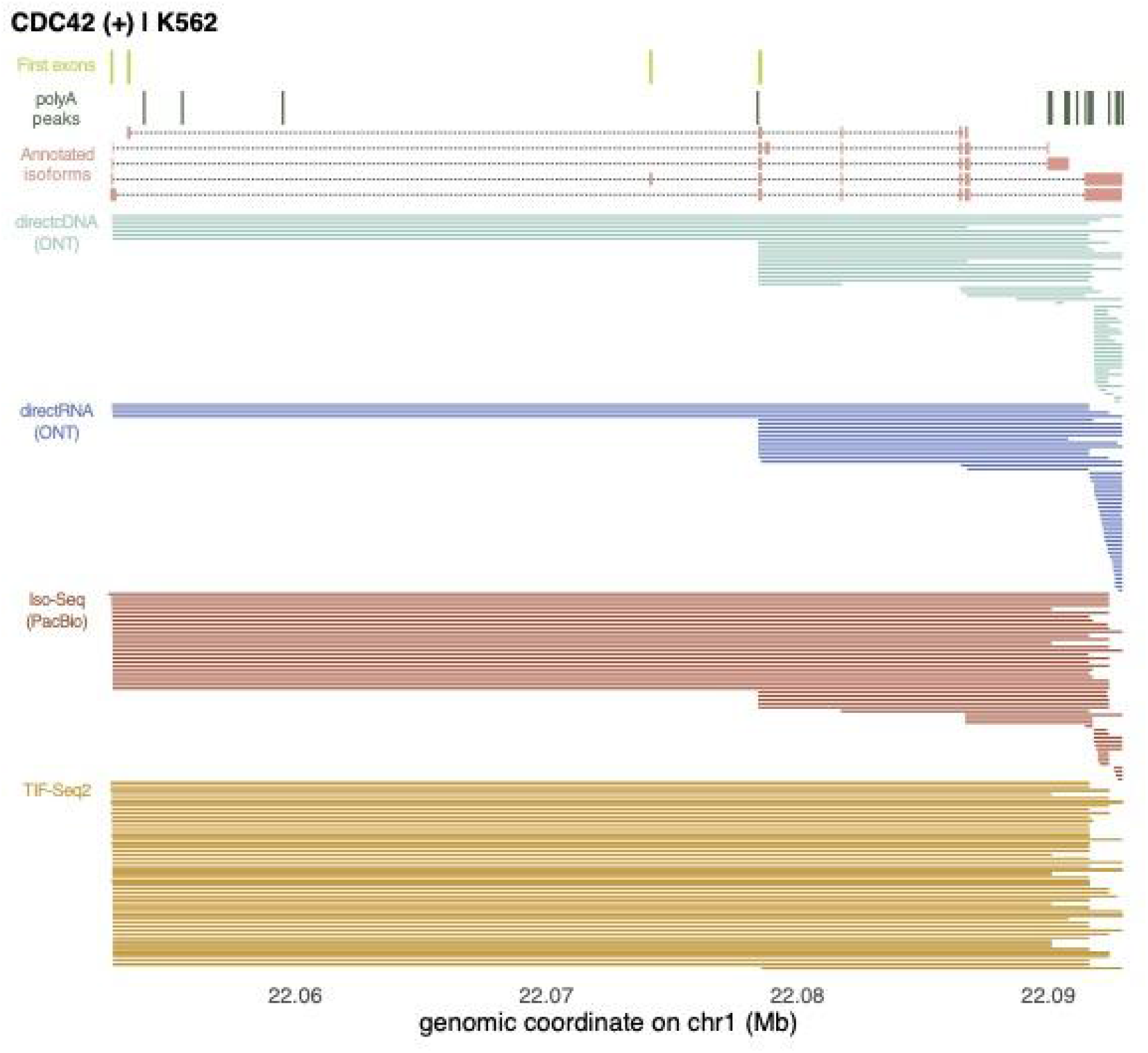
LRS reads show large variability in start and end coordinates across a gene. Representative example of long-read RNA sequencing reads in K562 cells for CDC42. To enhance visualization, we randomly sampled 50 reads from each sequencing technology. Each line shows the span between the first coordinate and the last coordinate of the read, for ONT direct cDNA (teal), ONT direct RNA (*blue*), and PacBio Iso-Seq (*red*). For TIF-Seq2 reads (*orange*), each line shows the span between the inferred start and ends from the circularized read. At the top of the figure are annotated genomics features of CDC42, including the HITindex first exons (*light green*, topmost track), polyA peaks (*dark green*, second track from the top), and annotated isoforms (*pink*, third track from the top).

First, LRS technologies show larger variability in terminal end positions than the orthogonal TIF-Seq2 short read approach. For instance, there are large variability number of transcript start sites identified across sequencing technologies for the CDC42 gene in K562 cells (**Figure 1**), with 5, 12, 5, and 1 starts site clusters supported by at least 2 reads starts that are less than 25nt away from each other across directcDNA, directRNA, Iso-Seq, and TIF-Seq2, respectively. There is less variability in transcript end site identification, with 3, 4, 3, and 3 end site clusters for CDC42 in K562 cells across directcDNA, directRNA, Iso-Seq, and TIF-Seq2, respectively. Notably, especially for read starts, many of the identified terminal positions do not overlap with terminal coordinates of annotated CDC42 isoforms or with empirically derived 5’ exon or 3’ polyA sites (**Figure 1**). While this trend is noticeable for all sequencing technologies, the variability in read start and end positions was lower for PacBio compared to both ONT protocols. For example, for each sequencing approach, 8%, 8%, 54%, and, 98% of reads had terminal ends that overlapped an annotated terminal end for directcDNA, directRNA, Iso-Seq, and TIF-Seq2, respectively.

Second, many LRS reads are derived from a single exon. For instance, 52%, 64%, 30%, and 0% of reads across directcDNA, directRNA, Iso-Seq, and TIF-Seq2, respectively, exhibited no splicing and were fully contained within the 3’ most exon of CDC42, despite there being no annotated single exon isoforms of CDC42. Together, these two features of LRS reads suggested the possibility of either widespread spurious terminal ends delineated by LRS reads or widespread novel isoform usage that was inconsistently detected by different sequencing approaches.

### The ends of LRS reads do not accurately recapitulate the annotated terminal ends of mRNA molecules

Based on our direct observation of short 3’ terminal reads for CDC42 (**Figure 1**), we first systematically assessed the extent of single exon reads sequencing methods. To do so, we measured the proportion of reads with their start and end positions within the boundaries of a single annotated exon (**Figure 2A**). Over 25% of reads generated by ONT methods can be classified as single exon reads, while Iso-Seq and TIF-Seq2 datasets have very low proportions of these single exon reads. These observations indicate that the presence of these reads represents technical differences between the protocols during library preparation or sequencing and lead to spurious transcript boundaries in reads.

**FIGURE 2:**
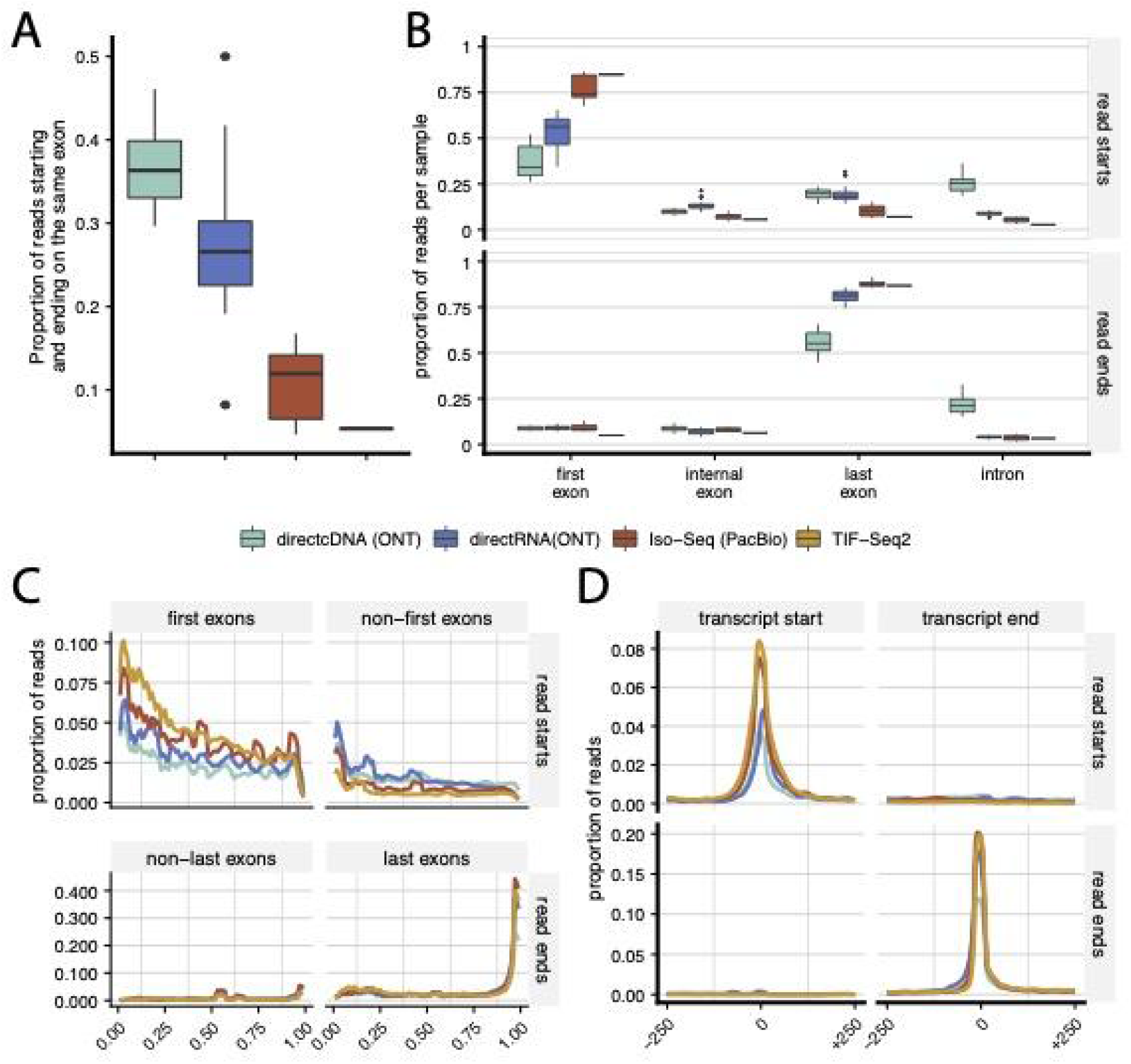
The ends of LRS reads do not accurately recapitulate the annotated terminal ends of mRNA molecules. **(A)** Proportion of reads that start and end within the same exonic feature. **(B)** The proportion of reads that start (*top*) or end (*bottom*) in HITindex classified first exons, internal exons, last exons, or annotated introns across data from 3 different LRS technologies and TIF-seq data. **(C)** The distribution of read starts (*top*) or ends (*bottom*) across terminal or non-terminal exons. The y-axis represents the proportion of reads per sample within each fractional position across the exonic feature, calculated using a sliding window of 0.01 across the feature. **(D)** The distribution of read starts (*top*) or ends (*bottom*) around annotated transcription start (*left*) or end (*right*) sites. The y-axis represents the proportion of reads per sample, calculated using a sliding window of 0.01 around the feature.

Next, we quantified the proportions of reads starting or ending in exons or introns, separating exons by their first, internal, or last position within the gene as determined by the HITindex (Methods, (28)). We find that a substantial proportion of read starts and ends do not fall within the expected first or last exon, respectively (**Figure 2B)**. This is especially true for read starts in the ONT approaches, in which more than 50% of the read starts in directcDNA libraries and more than 40% of read starts in directRNA libraries did not fall within first exons. This suggests that these read stars are aligned elsewhere in the gene and we find that for ONT methods, a higher proportion of read starts are found within the last exons. These results point to the presence of 5’ truncation in LRS reads, particularly in those from ONT approaches. Since there is a relatively low proportion of reads across approaches that fall in introns, these reads likely represent mature mRNA molecules and truncation of the 5’ end likely occurring during library preparation, sequencing, or processing of the data. The presence of truncation is also supported by the relatively shorter reads generated by directRNA and directcDNA approaches (**Figure S1, Table S1**).

When we look more specifically at the reads that started in the first exons, we see that a higher proportion of reads from the Iso-Seq and TIF-Seq2 methods start closer to the beginnings of first exons (as expected if they represent transcription start sites), while the precise start sites were more distributed across the first exon in both ONT methods (**Figure 2C**). For non-first exons (internal or last exons), we observed the opposite trend, where reads from ONT methods were more likely to start close to the beginning of these exons. In contrast, among reads that end in last exons, we see that the precise end sites were mostly found close to the end of the last exons, regardless of technology, with directcDNA reads tending to show slightly worse truncation even at the ends of the reads (**Figure 2C)**. When we conduct a similar analysis centered around annotated transcript start sites or end sites, we again see that Iso-Seq and TIF-Seq2 show the greatest proportions of read start and end sites located at the transcript start and end sites, respectively (**Figure 2D**). The distribution of read starts from ONT methods again show a slight shift towards positions downstream of the transcript start site. Finally, for all sequencing techniques other than directcDNA, the read ends are very closely distributed around the transcript end sites. Together, these results support systematic truncation of LRS reads, particularly at the 5’ end of reads, with more severe truncation occuring in reads from ONT approaches.

### Conditioning on annotated mRNA terminal exons retains reads likely to be full length transcripts

Given our observations of widespread truncation, especially at the 5’ end of reads, we devised a filtering strategy to differentiate truncated from full-length reads. To do so, we turned to databases of empirically derived transcription start and end sites. Specifically, we used HITindex first exons (28) and 5’ Cap Analysis of Gene Expression (CAGE) peaks (27) as annotations that likely enrich for true start sites and the polyA sites in the PolyASite 2.0 database (32) as annotations that likely enrich for true end sites. Using sites in these databases, we first assessed the proportions of reads that started and ended in first exons or CAGE peaks, and polyA peaks, respectively (**Figure 3A, Figure S2**). We observed that most of the Iso-Seq and TIF-Seq2 reads per gene started within a first exon or CAGE peak and ended within a polyA peak. In contrast, more than half of the reads per gene in the ONT methods did not match first exons, CAGE peaks, or polyA peaks. We find that the overlap with CAGE peaks is a lot more variable, with many reads starting and ending within last exons whose start and end sites overlap CAGE and polyA peaks, respectively. Given recent reports that 10-15% of CAGE peaks are located in human last exons (33–36) and putative evidence of re-capping of short 3’ fragments in the cytoplasm (32), we decided to only use HITindex first exons for downstream filtering.

**FIGURE 3:**
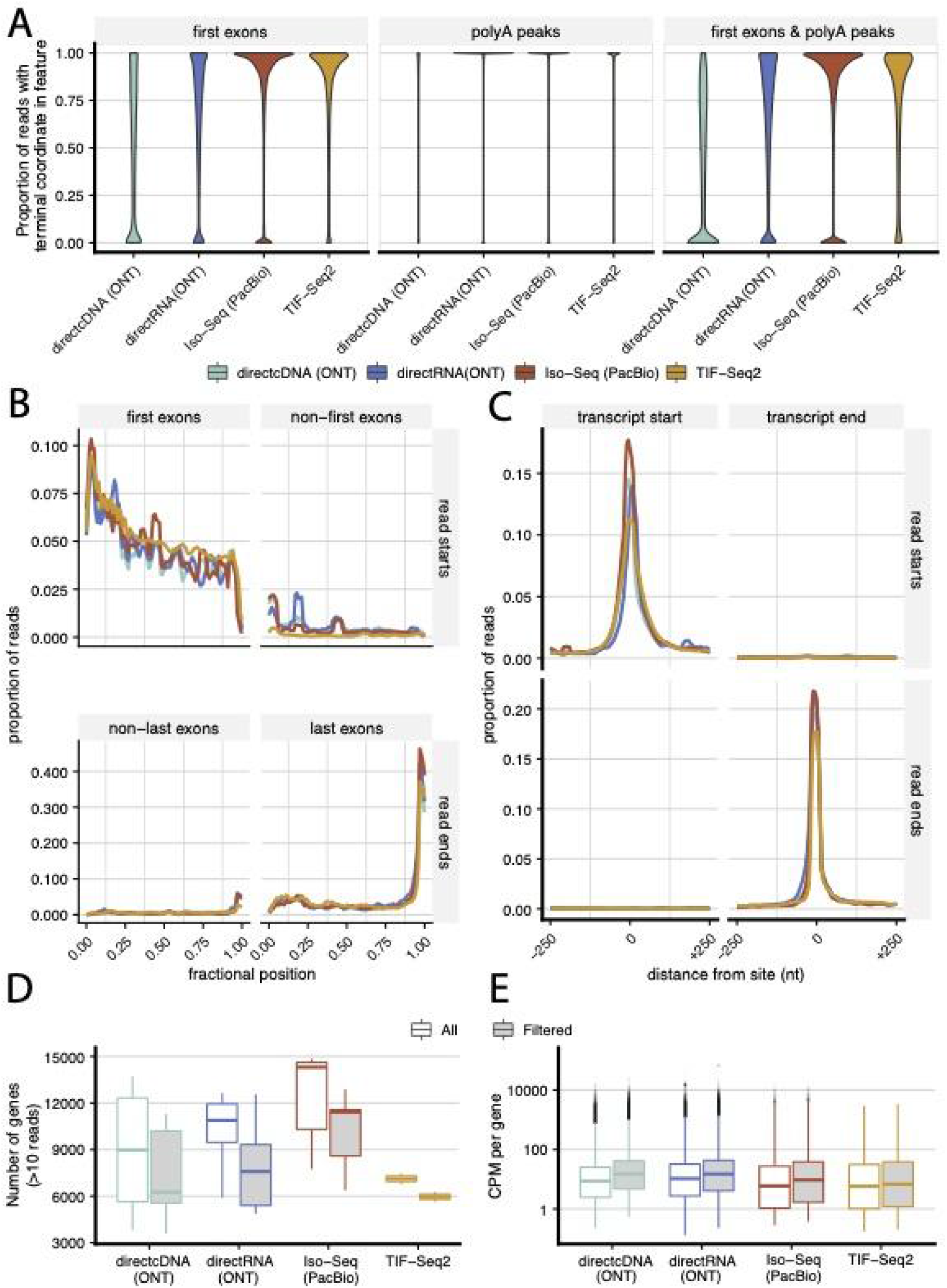
Conditioning on annotated mRNA terminal exons retains reads likely to be full length transcripts. **(A)** The distribution of the proportion of reads per gene starting in a first exon (left), ending in a polyA peak (*middle*), or meeting both of those criteria (*right*) for each of the 3 LRS technologies and TIF-seq data. **(B)** The distribution of read starts in first exons (*left*) or read ends in last exons (*right*) after conditioning on reads overlapping HIT-index and polyA annotations. The y-axis represents the proportion of reads per sample within each fractional position across the exonic feature, calculated using a sliding window of 0.01 across the feature. **(C)** The distribution of read starts (*top*) or ends (*bottom*) around annotated transcription start (*left*) or end (*right*) sites after conditioning on reads overlapping terminal annotations. The y-axis represents the proportion of reads per sample, calculated using a sliding window of 0.01 around the feature. **(D)** The number of genes per sample with at least 10 reads before (*left*) or after (*right*) conditioning on reads overlapping terminal annotations. **(E)** The distribution of counts per million (CPM) values across genes before (*left*) or after (*right*) conditioning on reads overlapping terminal annotations.

To identify reads whose start and end sites have a high confidence of matching true terminal ends, we removed reads that did not start and end within a first exon and a polyA peak, respectively. To evaluate the performance of this filtering, we again examined the distribution of read starts and ends within (a) first or non-first exons, (b) last or non-last exons, (c) annotated transcript start sites, and (d) annotated transcript end sites (**Figure 3B, 3C**). We see that, upon filtering, the start sites of reads across all methods are positioned at the beginnings of first exons and evenly around transcript start sites. Similarly, end sites of reads are positioned at the ends of last exons and evenly around transcript end sites. However, upon filtering, the number of genes with at least 10 reads greatly decreases across libraries from all sequencing approaches (**Figure 3D**), with ONT showing the largest reduction (average 63.6%, 39.2%, 19.3%, 16.3% decrease for directcDNA, directRNA, Iso-Seq, and TIF-Seq2, respectively). Reassuringly the relative gene expression levels (quantified by counts per million (CPM)) appear to be relatively consistent before and after filtering for each of the methods (**Figure 3E**). This lends credence to the strategy of filtering out reads that do not match empirically derived transcription start or end sites as a means of obtaining more confident terminal end counts while still allowing quantification of relative gene expression levels.

## DISCUSSION

In this study, we systematically characterized the accuracy of terminal ends in reads from long-read mRNA ONT and PacBio data. We first observed that reads from all sequencing technologies show a large range of start and end coordinates within genes. While we cannot absolutely rule out the possibility that many of these reads arise from novel isoforms or terminal ends, we see evidence that most of these reads arose from technical issues, the extent of which varied according to the library preparation and sequencing approach. Moreover, the start or end coordinates of these reads often did not match either annotated or empirically derived 5’ or 3’ ends of mRNA molecules. Our results showed pervasive 5’ truncation in these data, particularly for ONT data types, which may lead to the spurious identification of transcription start sites or first exons. The effects of these truncation errors affected both the number of and quantification of genes that could be analyzed and underscore the need for caution when using LRS data to identify and quantify terminal ends.

While filtering these reads to only retain reads whose terminal ends are supported by empirically derived 5’ or 3’ mRNA ends increased our confidence in these biological findings by reducing technical errors, these steps greatly reduced the number of reads. Filtering using either first exons or CAGE peaks seems to perform similarly to reduce spurious terminal ends, however, using the latter dataset carries forward short or truncated reads that start and end in last exons and are likely to be spurious. While it is possible that these CAGE peaks represent biologically accurate capped RNAs, they may not result from PolII transcribed mRNAs (33), highlighting the issues with using databases that broadly capture functional marks. Thus, using CAGE peaks to filter may lead to erroneous identification of mRNA isoforms.

One advantage of LRS is the ability to discover novel isoforms; however, our observations illustrate the challenge in differentiating true novel observations of terminal site usage from technical artifacts. While solutions to the low read accuracy for other parts of the genome have been proposed (22, 37), many of these solutions require additional third-party reagents or introduce concerns regarding potential biases and lack of efficiency. Moreover, read truncation is still present, and the analysis of these data also requires filtration and the loss of a substantial proportion of reads (22, 37).

These issues might be caused by both RNA quality, library preparation, sequencing, or analytical effects. These effects may start before or during library construction, leading to *in-vitro* truncation or degradation of RNA. In protocols calling for reverse transcription (RT), which include the use of a first-strand synthesis to form a structural splint for direct RNA sequencing in ONT and full cDNA synthesis for both the directcDNA ONT and PacBio IsoSeq approaches, limited processivity or limitations in RT enzymes may lead to truncation(38, 39)]. During library preparation, in-vitro truncation or degradation might occur during adapter ligation or other steps. In ONT sequencing, the characteristics of the pores themselves (*ie*. translocation speed too rapid to capture the last 10-15 nt (40)), spikes in electrical current or motor protein stalling that cause software to stop base calling (20, 41), or other software issues may also contribute to the observation of substantial read truncation. Rapid development of library preparation approaches, sequencing chemistry, and improvements in analysis algorithms may reduce these error rates.

Thus, although LRS continues to offer new possibilities for finer-scale resolution of essential biological and genomic processes, it is important to be cautious when selecting the specific application and interpreting data, as technical and analytical biases may affect the results and/or lead to misleading findings. We find that this risk is especially prevalent for studies intending to discover or quantify ends of mRNA transcripts. While filtration and other methods can improve confidence in these observations, these steps come at the cost of a greatly reduced number of reads and the reliance on external (often not sample matched) datasets reduces the ability to truly make novel observations.

## DATA AVAILABILITY

The script used to calculate truncation metrics can be found at https://github.com/ezecalvo/tss_tes_terminal_truncation. All publicly available used data is listed on Table S1.

## SUPPLEMENTARY DATA

## AUTHOR CONTRIBUTIONS

ECR: Conceptualization, Methodology, Formal analysis, Investigation, Writing - Original Draft, Visualization. RFD: Conceptualization, Investigation, Writing - Original Draft. AAP: Conceptualization, Investigation, Writing - Review & Editing, Supervision.

## ACKNOWLEDGEMENTS

We thank Ana Fiszbein, Christine Carroll, Zach Wakefield, Steven T. Mick, and the Pai Lab for help with databases, discussions and feedback on the manuscript.

## FIGURE LEGENDS

**Figure S1:**
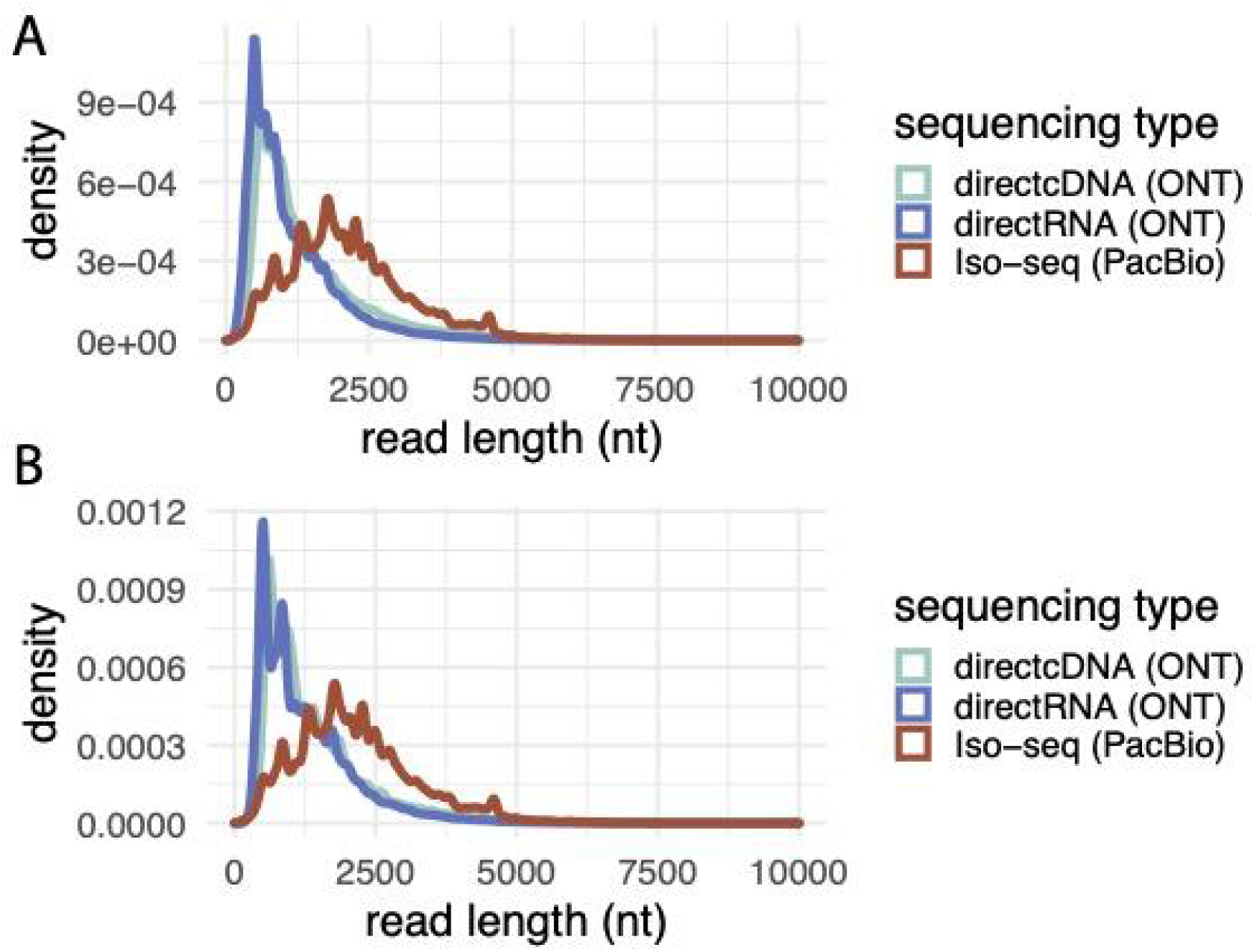
Distribution of read lengths for long-read RNA sequencing technologies before (A) and after (B) terminal end filtering steps.

**Figure S2:**
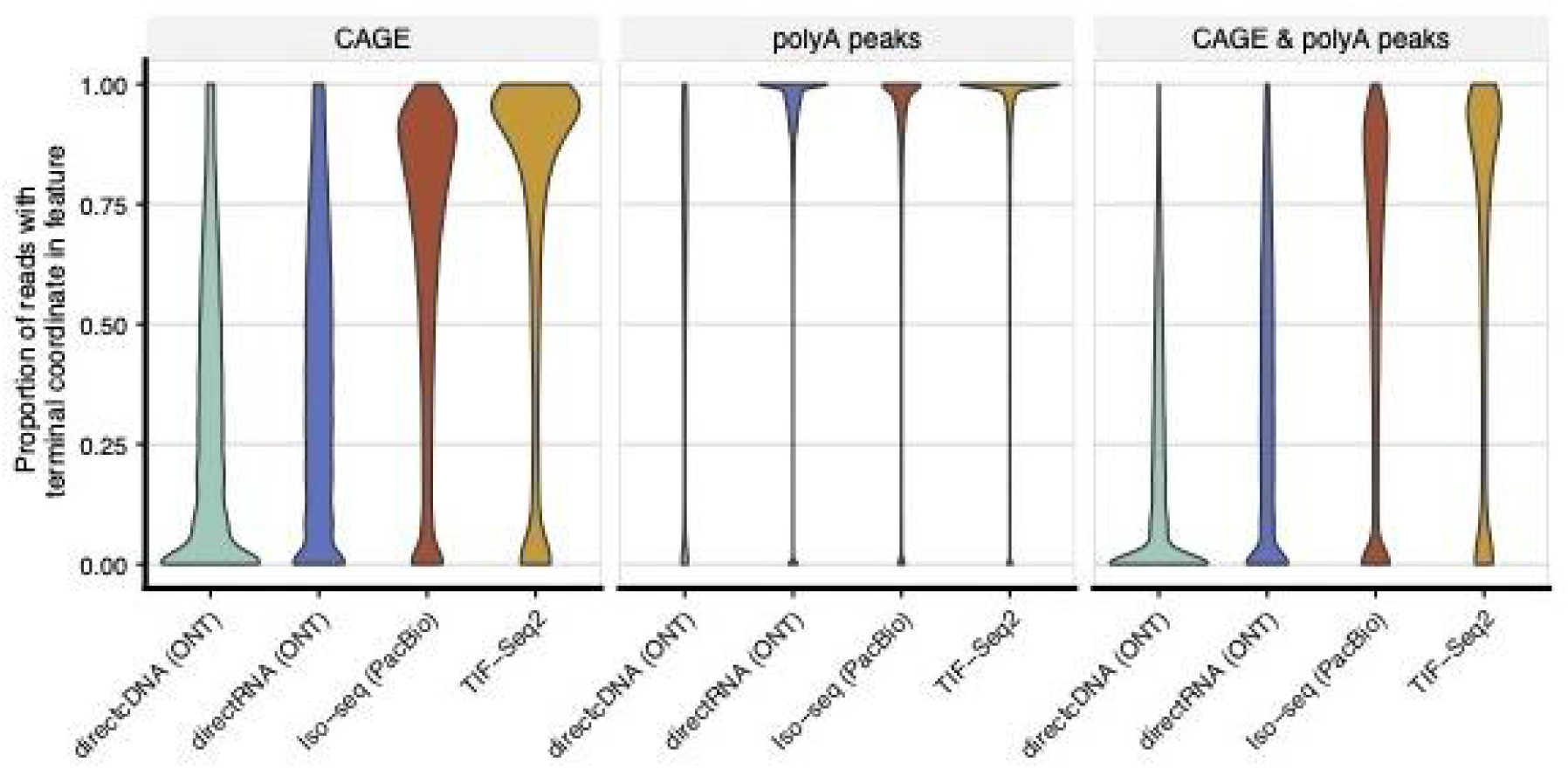
Conditioning on long-read RNA sequencing reads aligning with CAGE peaks. The distributions of the proportion of reads per gene starting in a CAGE peak (*left*), ending in a polyA peak (*middle*), or meeting both of those criteria (*right*) for each of the three LRS technologies and TIF-seq data are shown.

